# MetaWRAP - a flexible pipeline for genome-resolved metagenomic data analysis

**DOI:** 10.1101/277442

**Authors:** Gherman V Uritskiy, Jocelyne DiRuggiero, James Taylor

## Abstract

**Background:** The study of microbiomes using whole-metagenome shotgun sequencing enables the analysis of uncultivated microbial populations that may have important roles in their environments. Extracting individual draft genomes (bins) facilitates metagenomic analysis at the single genome level. Software and pipelines for such analysis have become diverse and sophisticated, resulting in a significant burden for biologists to access and use them. Furthermore, while bin extraction algorithms are rapidly improving, there is still a lack of tools for their evaluation and visualization.

**Results:** To address these challenges, we present metaWRAP, a modular pipeline software for shotgun metagenomic data analysis. MetaWRAP deploys state-of-the-art software to handle metagenomic data processing starting from raw sequencing reads and ending in metagenomic bins and their analysis. MetaWRAP is flexible enough to give investigators control over the analysis, while still being easy-to-install and easy-to-use. It includes hybrid algorithms that leverage the strengths of a variety of software to extract and refine high-quality bins from metagenomic data through bin consolidation and reassembly. MetaWRAP’s hybrid bin extraction algorithm outperforms individual binning approaches and other bin consolidation programs in both synthetic and real datasets. Finally, metaWRAP comes with numerous modules for the analysis of metagenomic bins, including taxonomy assignment, abundance estimation, functional annotation, and visualization.

**Conclusions:** MetaWRAP is an easy-to-use modular pipeline that automates the core tasks in metagenomic analysis, while contributing significant improvements to the extraction and interpretation of high-quality metagenomic bins. The bin refinement and reassembly modules of metaWRAP consistently outperform other binning approaches. Each module of metaWRAP is also a standalone component, making it a flexible and versatile tool for tackling metagenomic shotgun sequencing data. MetaWRAP is open-source software available at https://github.com/bxlab/metaWRAP.

## Background

The study of microbial communities through whole metagenomic (WMG) shotgun sequencing opens new avenues for the investigation of the metabolic potentials of microbiomes, in addition to their taxonomic composition [1-3]. This greatly improves the ability to interpret and predict functional interactions, antibiotic resistance, and population dynamics of microbiomes, with applications in human health, waste treatment, agriculture, and environmental stewardship [4-6]. WMG shotgun sequencing reads from hundreds to thousands of community members generates unique challenges for data analysis and interpretation [3, 7]. Software for WMG data analysis have grown in number and complexity, improving our ability to process, analyze, and interpret such data [8-12]. However, these tools are burdensome for biologists to work with. As the field of WMG expands, comprehensive and accessible software for unified analysis of metagenomic data is needed [7, 11].

Running a WMG analysis requires investigators to find the best currently available tools, install and configure them on a cluster, address conflicting libraries and environmental variables, and write scripts to convert outputs from one tool into the correct format to input into the next tool [13, 14]. These challenges present a major burden to anyone attempting metagenomic analysis, especially for investigators without computational experience, hindering progress of microbial genomics as a field [15]. Existing automated pipelines and cloud services lack modularity, do not give users control over the analysis, and often lack functions for genome-resolved metagenomics, the extraction of putative genomes (bins) through the binning of metagenomic assemblies [14, 16-19].

Genome-resolved metagenomics allows for metabolic reconstruction of individual taxa and microbiome comparison at a finer scale. While a number of sophisticated tools such as CONCOCT, MaxBin, and metaBAT have been developed to address binning, it is still an actively improving field [9, 20-22]. Qualities of a metagenomic bin are (1) completion, the level of coverage of a population genome, and (2) contamination the amount of sequence that do not belong to this population from another genome. These metrics can be estimated by counting universal single-copy genes within each bin [23, 24]. CheckM improves on this by checking for single-copy genes that a genome of the bin’s taxonomy is expected to have [25].

Most metagenomic binning tools extract bins by clustering together scaffolds that have similar sequence properties, such as K-mer composition and codon usage, and similar read coverages across multiple samples [26, 27]. Because no single binning approach is superior in every case, bin consolidation tools attempt to combine the strengths and minimize the weaknesses of different approaches. DAS_Tool predicts single-copy genes in all the provided bin, aggregates bins with overlapping genes, and extracts a more complete consensus bin from each aggregate[28]. This collapsing approach significantly improves the completion of the bins. Binning_refiner, on the other hand, splits the contigs into bins such that all contig division boundaries of the original predicted bins are satisfied. This breaks the contigs into many more bins, reducing contamination[29]. Both of these approaches consolidate sets of bins from different methods and result in a superior bin set, but they have limitations – DAS_Tool increases completion at the expense of introducing contamination, while Binning_refiner prioritizes purity, but loses completeness. Another way to improve draft genome quality that is relatively unexplored is bin reassembly – extracting reads that belong to a given bin and assembling them separately from the rest of the metagenome. With proper benchmarking, this approach could significantly improve the quality and downstream functional annotation of at least some bins in a microbial community.

Because the field of shotgun metagenomics is relatively new, there is a lack of software to inspect, analyze, and visualize metagenomic bins. While there are tools that can accurately predict the taxonomy of metagenomic scaffolds (such as Taxator-tk), there is no tool to classify entire metagenomic bins[30, 31]. Similarly, there are many ways to estimate the coverage of scaffolds based on read alignment depth, but no way to find the coverages of entire bins across many samples[32, 33]. Finally, there no tool to visualize draft genomes in context of whole metagenomic communities. The need for an easy-to-use integrated tool for WMG data analysis, as well as the lack of available tools for metagenomic bin analysis motivated the construction of MetaWRAP.

## Implementation

MetaWRAP is command line software for Unix-based systems that calls on a collection of modules, each being a standalone program addressing one aspect of WMG data processing or analysis (Figure 1). Each module is a shell script pipeline that takes in a variety of input files parameters through command line flags. The modules call upon numerous installed software as well as custom Python 2.7 scripts. MetaWRAP relies on the modules folder (metawrap-modules), the scripts folder (metaWRAP-scripts), and a file containing paths to databases (config-metawrap) to be available in the PATH (see Methods in Additional file 1). MetaWRAP is hosted on github (https://github.com/bxlab/metaWRAP), distributed through Anaconda[34], and can be easily installed locally and on remote clusters. The metawrap-mg conda package (https://anaconda.org/ursky/metawrap-mg) includes metaWRAP and the necessary software for running any metaWRAP modules. The databases required by some modules need to be downloaded and unpackaged as described in the metaWRAP database download guide, and their paths indicated in the config-metawrap file. MetaWRAP v0.7 was used in all benchmarking runs.

**Figure 1:**
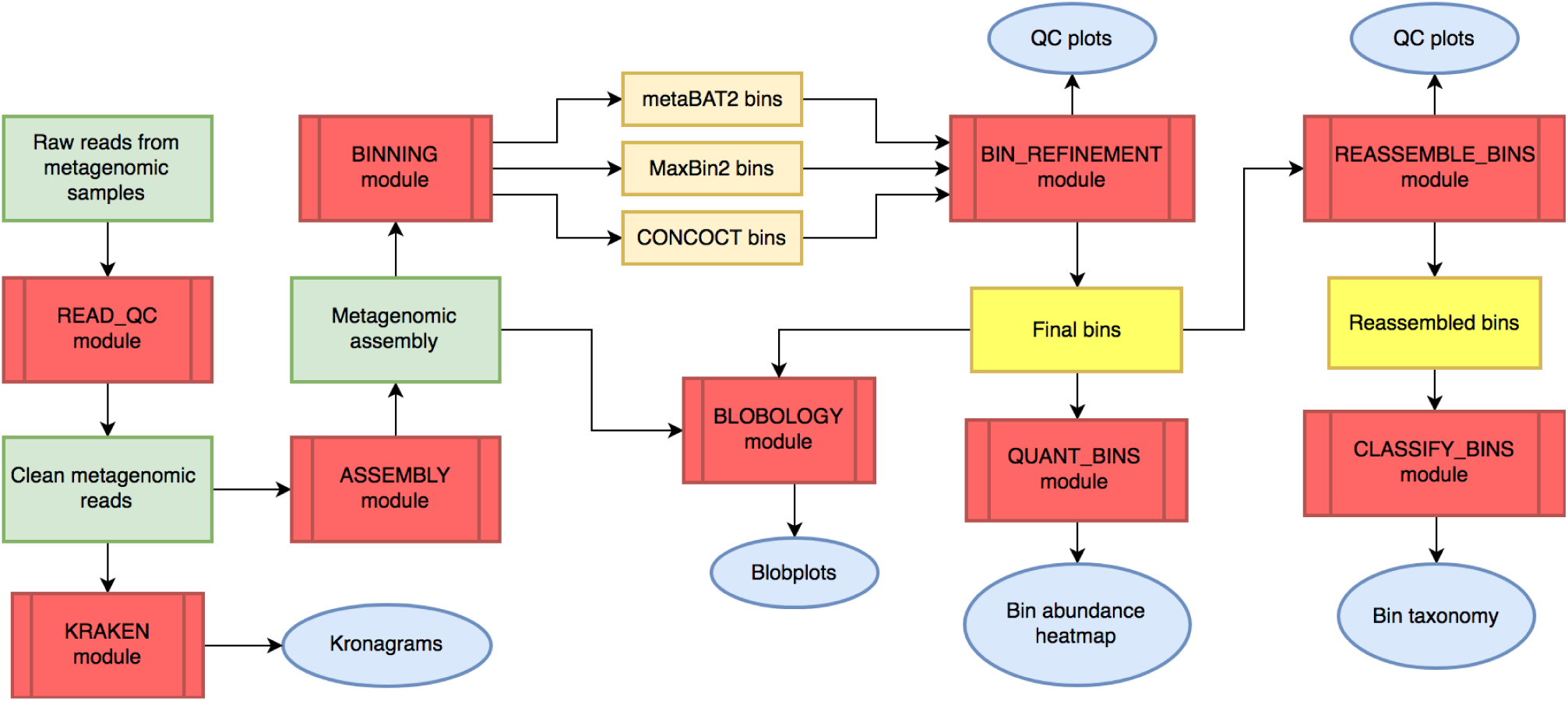
Overall workflow of metaWRAP. The flowchart illustrates the main interactions between the modules (red), metagenomic data (green), intermediate (orange) and final bin sets (yellow), and data reports and figures (blue).

The metaWRAP installation produces a bioinformatics environment with over 150 commonly used bioinformatics software and libraries (Additional file 2: Figure S1). MetaWRAP itself is a collection of modules, each of which uses a variety of pre-existing and newly developed software and databases to accomplish a specific step of metagenomic analysis. Unlike existing metagenomic wrappers and cloud services, metaWRAP retains modularity and grants the user control of the analysis pipeline. The user may follow the intuitive workflow starting from raw metagenomic shotgun sequencing reads all the way to high-quality draft genomes and their functional annotation, or use only specific functions, as each module is also a standalone program

## Results and Discussion

### MetaWRAP is a flexible, modular pipeline

The metaWRAP installation produces a bioinformatics environment with over 150 commonly used bioinformatics software and libraries (Additional file 2: Figure S1). MetaWRAP itself is a collection of modules, each of which uses a variety of pre-existing and newly developed software and databases to accomplish a specific step of metagenomic analysis. Unlike existing metagenomic wrappers and cloud services, metaWRAP retains modularity and grants the user control of the analysis pipeline. The user may follow the intuitive workflow starting from raw metagenomic shotgun sequencing reads all the way to high-quality draft genomes and their functional annotation, or use only specific functions, as each module is also a standalone program (Figure 1).

The first metaWRAP module, the metaWRAP-Read_qc module, trims the raw sequence reads, removes human contamination, and produced quality reports for each of the sequenced samples. The reads from all given samples can be then assembled with the metaWRAP-Assembly module using MegaHit[35] or metaSPAdes[36], which also produces an assembly report. Both the reads from each sample and the assembly can be taxonomically profiled with the Kraken[30] module, producing interactive kronagrams[37] of community taxonomy. The assembly is then binned with the metaWRAP-Binning module by three metagenomic binning software – MaxBin2, metaBAT2, and CONCOCT[19, 21, 22].

The other modules of metaWRAP focus on refining, analyzing, and visualizing metagenomic bins from either the Binning module or other sources. MetaWRAP-Bin_refinement module hybridizes to three bin sets with Binning_re-finer[29], and then finds the best version of each bin based on completion and contamination metrics estimated with CheckM[25] (Additional file 3: Figure S2). The scaffolds in the final bin set are then de-replicated, and a report of their completion, contamination, and other metrics is produced. MetaWRAP-Reassemble_bins can then be used to reassemble the reads belonging to each bin, improving their N50, completion, and contamination (Additional file 4: Figure S3). The resulting bins can be visualized by using the metaWRAP-Blobology module[38], which plots the contigs of the joint assembly on a blob plot, annotating them with their taxonomy and bin membership. The metaWRAP-Quant_bins module can be used to quickly estimate the abundance of each bin in each of the metagenomic samples. MetaWRAP-Classify_bins can be used to conservatively, but accurately estimate their taxonomy. Finally, the bins can be functionally annotated with the metaWRAP-Annotate_bins module.

### Compute time of metaWRAP modules

The runtime of each metaWRAP module was evaluated on a subset of the Human Intestinal Tract (MetaHIT) survey [39]. The same subset is used in the metaWRAP tutorial page on GitHub. The data contained 3 WMG samples, totaling 145.8 million 75bp paired-end reads, or 21.9Gbp of sequencing data. MetaWRAP was used to analyze this dataset on a Linux server with 24 cores and 100GB of RAM. All modules were run on default settings, and the total runtime of each module was recorded (Additional file 5: Module runtime). The entire pipeline was completed in 5h36m, with the majority of compute time dedicated to the Read_qc, Binning, Bin_refinement, and Reassemble_bins modules. With the exception of CONCOCT [19], the programs wrapped into metaWRAP can take advantage of multi-core systems, and scale well with larger datasets. MetaWRAP itself also parallelizes processes when possible.

### MetaWRAP-Bin_refinement improved bin predictions in synthetic data

To test the efficacy of the metaWRAP-Bin_refinement module at consolidating and improving bin sets, we used synthetic metagenomic data sets of varying complexity from the Critical Assessment of Metagenomic Interpretation (CAMI) study[9]. The “gold standard” assemblies from the “high”, “medium”, and “low” diversity challenges were first binned with metaBAT2, Maxbin2, and CONCOCT[19, 21, 22] using the metaWRAP-Binning module, and the resulting three bin sets were then consolidated with DAS_Tool[28], Binning_refiner[29], and metaWRAP-Bin_refinement. The completion and contamination of the bins in the original and refined bin sets were evaluated with CheckM[25] (Additional file 6: Figure S4) and Amber[40] (Additional file 7: Figure S5). True recall and precision for each bin calculated with Amber were converted to completion and contamination percentages to be comparable to the CheckM results (Figure 2). We found that metaBAT2 consistently outperformed MaxBin2 and CONCOCT, producing a total of 385 high quality bins between all the challenges (completion greater than 90% and contamination less than 5%), and 271 near-perfect bins (completion greater than 95% and contamination less than 1%). MaxBin2 came in second with 275 high quality bins and 164 near-perfect bins. CONCOCT performed rather poorly in all but the smallest CAMI challenge data sets, producing 58 high quality bins and 40 near-perfect bins.

**Figure 2:**
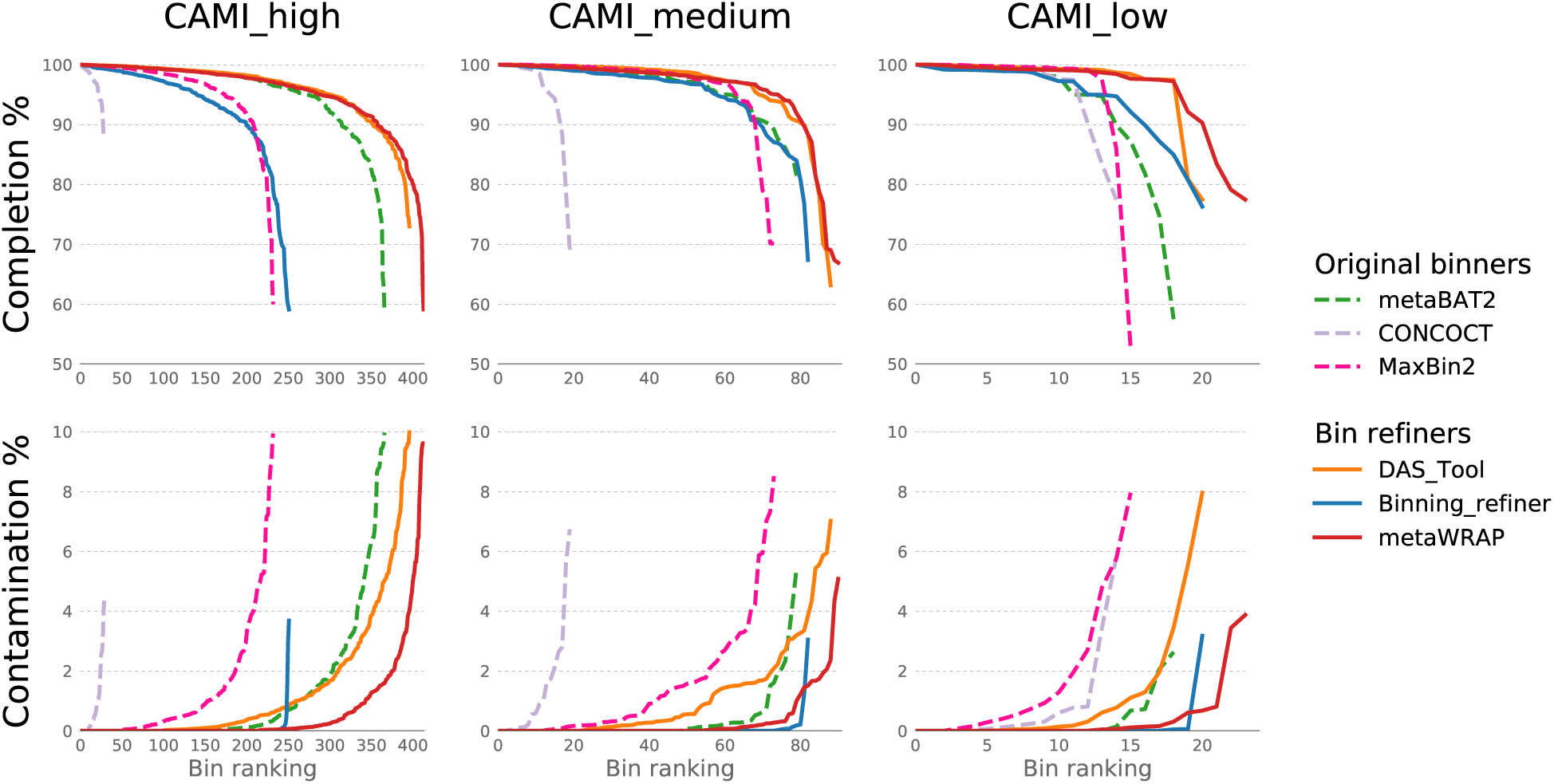
True completion and contamination of bins recovered from the CAMI’s high, medium, and low complexity synthetic data sets using original binning software (metaBAT2, MaxBin2, and CONCOCT) and software consolidating the original sets (DAS_Tool, Binning_refiner, and metaWRAP’s Bin_refinement module). Only bins with≥50% completion and≤10% contamination are shown.

In the consolidated bin sets, DAS_Tool produced 426 high quality bins and 263 near-perfect bins across all CAMI challenges, while Binning_refiner produced 289 and 210 bins, respectively. DAS_Tool consistently produced high completion bins, however these bins had relatively high contamination, which is a result of the aggregation approach that DAS_Tool takes. Binning_refiner on the other hand produced very pure bins with its splitting approach, however it did so at the expense of significantly reduced completion. MetaWRAP-Bin_refinement produced bins that had both high completion and low contamination. In total, it produced 457 high quality bins and 339 near-perfect bins (Figure 2) due to both a splitting and aggregation step. These results confirmed that metaWRAP not only consistently improved bin sets through its consolidation approach, but it also outperformed other consolidation algorithms in data sets of varying complexity.

The use of CheckM (Additional file 6: Figure S4) and Amber (Figure 2) to evaluate the binning sets produced similar results, although overall CheckM slightly overestimated both completion and contamination of the produced bins. More importantly, the relative performance of the six binning approaches was the same when evaluating with CheckM or Amber. This validated the use of CheckM for benchmarking binning results in data sets where the true genomes remain unknown.

### Bin_refinement improved bin predictions in real data

MetaWRAP was also benchmarked against real metagenomic data, using water, gut, and soil microbiome WGS data sets. The water data set was from a brackish water survey of the Baltic Sea [41] and included 36 samples for a total of 196Gbp of sequence. The gut data set came from the Metagenomic of the Human Intestinal Tract (MetaHIT) survey [39] and consisted of 50 samples and a total of 144Gbp. The soil data set was from grassland soil microbial communities from Angelo Coastal Reserve [28] and included 6 samples for a total of 481Gbp of sequencing data.

Samples from each microbiome type were pre-processed through the metaWRAP-Read_qc module to trim reads and remove human contamination, and the Kraken module was used to obtain the taxonomic profile of the communities (Additional file 8: Community taxonomy). The water samples were dominated by Alphaproteobacteria and Actinobacteria, the gut samples were dominated by Bacteroidetes and Clostridia, and the soil samples comprised of a wide variety of Proteobacteria and Terrabacteria (Figure 2).

The quality-controlled reads were then co-assembled with the metaWRAP-Assembly module and the assemblies binned with metaBAT2 Maxbin2, and CONCOCT using the metaWRAP-Binning module. The resulting three bin sets for each microbiome type were consolidated with DAS_Tool, Binning_refiner, and metaWRAP-Bin_refinement, and the completion and contamination of the resulting bins were evaluated with CheckM (Figure 4). Across the original binning software, metaBAT2 consistently produced the best sets of bins when compared to MaxBin2 and CONCOCT, with 202, 146, and 88 acceptable quality bins (comp≥50%, cont≤10%) in water, gut, and soil samples, respectively. MaxBin2 had 151, 98, and 40 bins, and CONCOCT 65, 121, and 39 bins. Despite incorporating all the binning methods, DAS_Tool was unable to improve the original bin sets, producing 198, 130, and 63 acceptable quality bins in water, gut, and soil samples, respectively. DAS_Tool performed relatively well at higher bin completion ranges (≥80%), although at the expense of increased contamination. Binning_refiner performed similarly, with 206, 138, and 83 bins in water, gut, and soil data sets, respectively. The bins from Binning_refiner were less complete, but also had significantly lower contamination than bins in the original bin sets. MetaWRAP’s Bin_refinement module produced 235, 175, and 134 acceptable quality bins in water, gut, and soil samples, respectively, significantly outperforming all other tested approaches. The module uses Binning_refiner in its pipeline to hybridize the input bin sets, and then choses the best version of each bin from the original and hybridized sets. Because the Bin_refinement module leverages the strength of Binning_refiner but still has a collapsing step similar to DAS_Tool, it is able to match DAS_Tool’s high completion rankings, while retaining the low contamination rankings of Binning_refiner. Overall, MetaWRAP consistently produced the highest quality bin sets in all the tested metagenomic data sets, which ranged greatly in diversity, taxonomic composition, and sequencing depths.

It is important to note that the use of metaWRAP’s Bin_refinement module to improve binning predictions is not limited to the bin sets produced from the metaWRAP-Binning module (metaBAT2, MaxBin2, and CONCOCT). Bin sets from any 2 or 3 binning software may be used as input for the module. Furthermore, because the algorithm leverages the differences between the input bin predictions, it is also possible to use bin sets produced from different parameters of the same software as input.

### Bin_refinement adjusts to the desired bin quality

To consolidate the original and hybridized bin sets, metaWRAP-Bin_refinement chooses the best version of each bin based on their completion and contamination values. However, this selection is subjective, and depends on what the user believes to be the “best bin”. The minimum completion (-c) and maximum contamination (-x) options are key parameters that greatly alter the quality of the bins produced, as the module will dynamically adjust its algorithms to produce the maximum number of bins in this range.

To demonstrate the effects of changing the –c and –x parameters of metaWRAP’s Bin_refinement module, we ran the original bin sets from water, gut, and soil data sets with varying minimum completion (but fixed maximum contamination) (Additional file 9: Figure S6), and varying maximum contamination (but fixed minimum completion) (Additional file 10: Figure S7) parameters. When compared to the original Bin_refinement run (-c 50 –x 10), the module produced a greater number of bins at any given threshold when it was given custom –c and –x parameters. The improvements were especially noticeable at higher completion and lower contamination ranges. For example, MetaWRAP-Bin_refinement –c 90 –x 10 recovered 19, 18, and 1 (water, gut, and soil, respectively) extra bins with a minimum completion of 90%, when compared to the baseline –c 50 –x 10 run. Similarly, MetaWRAP-Bin_refinement with –c 50 –x 1 parameters extracted 8, 21, and 4 (water, gut, and soil, respectively) more bins at a maximum contamination of 1%, when compared to the baseline run. Unlike arbitrary and sometime confusing thresholding parameters in many other software, the minimum completion and maximum contamination options offer the user an intuitive way to parameterize the metaWRAP’s Bin_refinement module to their needs. This leads to significant increases in the number of quality bins they are able to extract from their data.

### Reassemble_bins significantly improved bin quality

MetaWRAP’s Reassemble_bins module improves a given set of bins through individual reassembly with SPAdes[42]. The module only replaces the original bins if the reassembled ones are better in terms of completion and contamination. Like the Bin_refinement module, the Reassemble_bins module takes in minimum completion (-c) and maximum contamination (-x) parameters to allow the user to define what they consider a “good” bin. The bins produced from the water, gut, and soil data with metaWRAP-Bin_refinement module runs (–c 50 –x 10) were run through the metaWRAP-Reassemble_bins module (-c 50 –x 10), and the resulting bins were re-evaluated with CheckM[25]. The Reassemble_bins module improved upon 78%, 98%, and 2% of the bins in the water, gut, and soil bin sets, respectively. The module significantly improved the water and bin set overall metrics, increasing their N50 and completion scores. Even more strikingly, the reassembly process significantly reduced contamination in these bin sets. (Figure 5). The success of the bin reassembly algorithm relies heavily on accurate and specific recruitment of the correct reads to each bin. In very diverse and heterogeneous communities such as those found in soil, the read recruitment may not be specific enough. This confused the assembler during the re-assembly stage, and resulted in an improvement for only a small fraction of the bins. However, draft genomes from gut and water samples were still significantly improved with the Reassemble_bins module despite their complexity (Figure 3).

**Figure 3:**
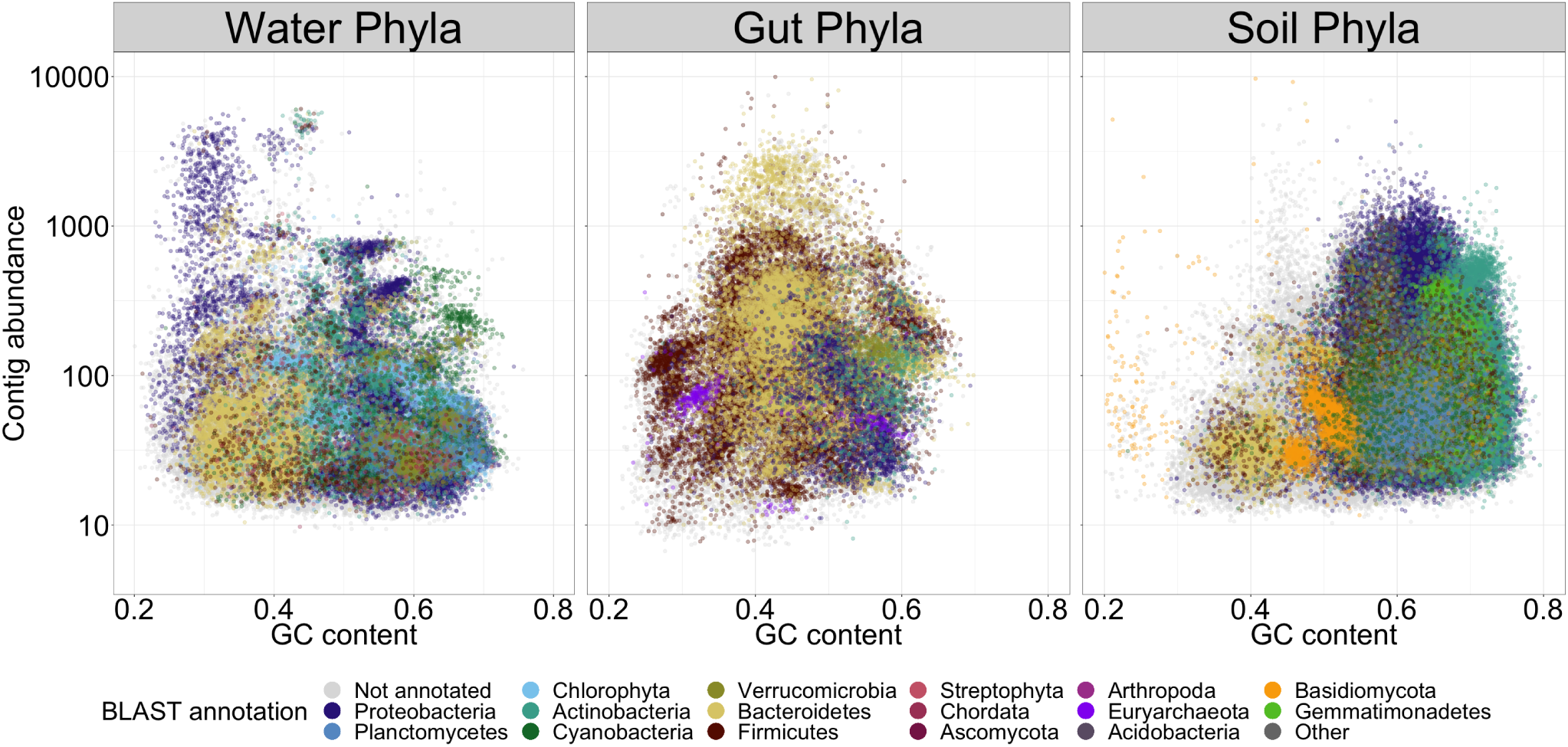
GC vs. abundance plots of contigs from water, gut, and soil metagenomes, produced with the Blobology module. Abundance of contigs was calculated from standardized read coverage in each sample. Contigs were annotated with their phylum taxonomy, as determined by BLAST

**Figure 4:**
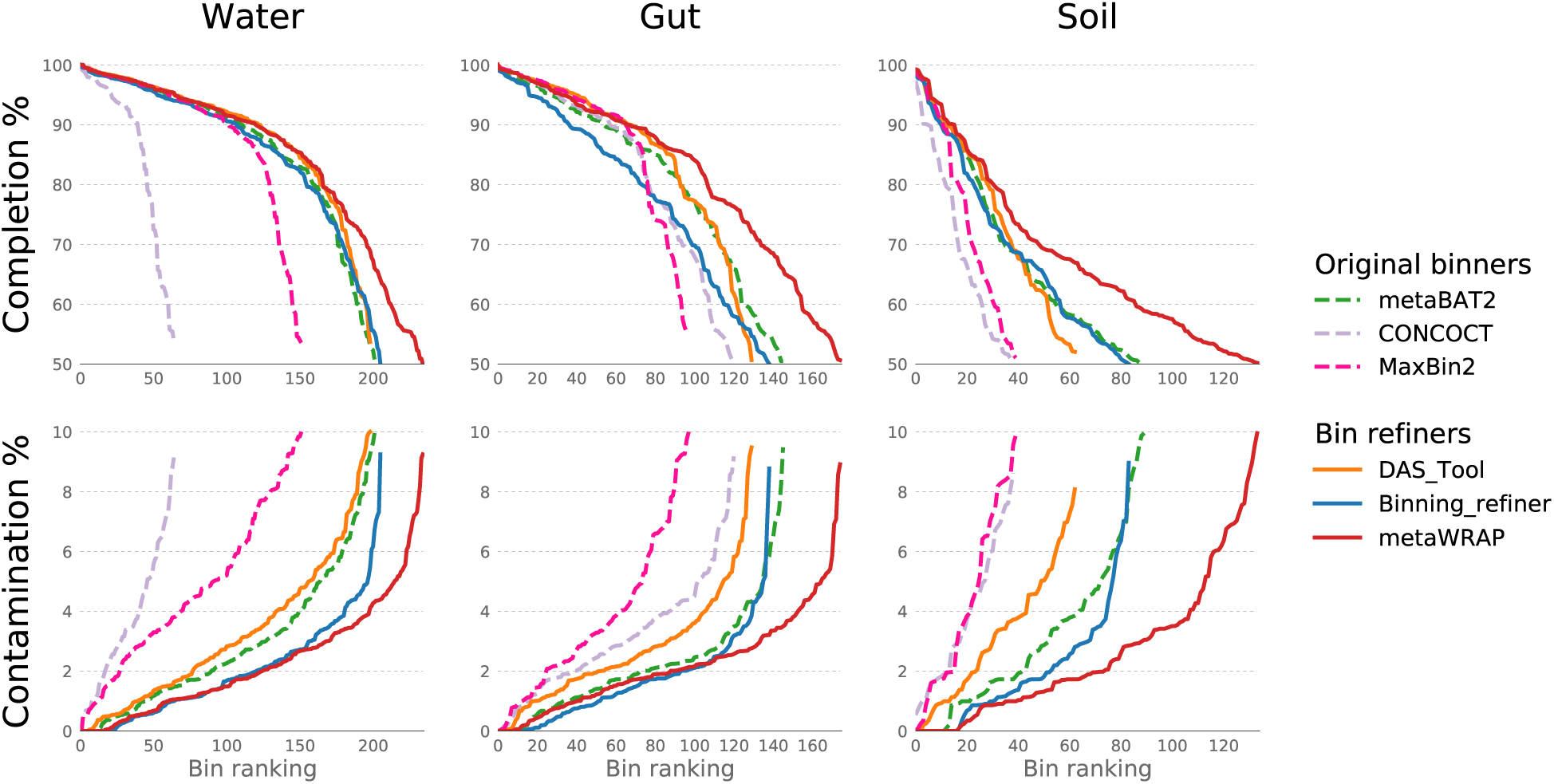
Completion and contamination of bins recovered from water, gut, and soil metagenomes using original binning software (metaBAT2, MaxBin2, and CONCOCT) and software consolidating the original sets (DAS_Tool, Binning_refiner, and metaWRAP’s Bin_refinement module). Only bins with≥50% completion and≤10% contamination are shown (estimated by CheckM).

**Figure 5:**
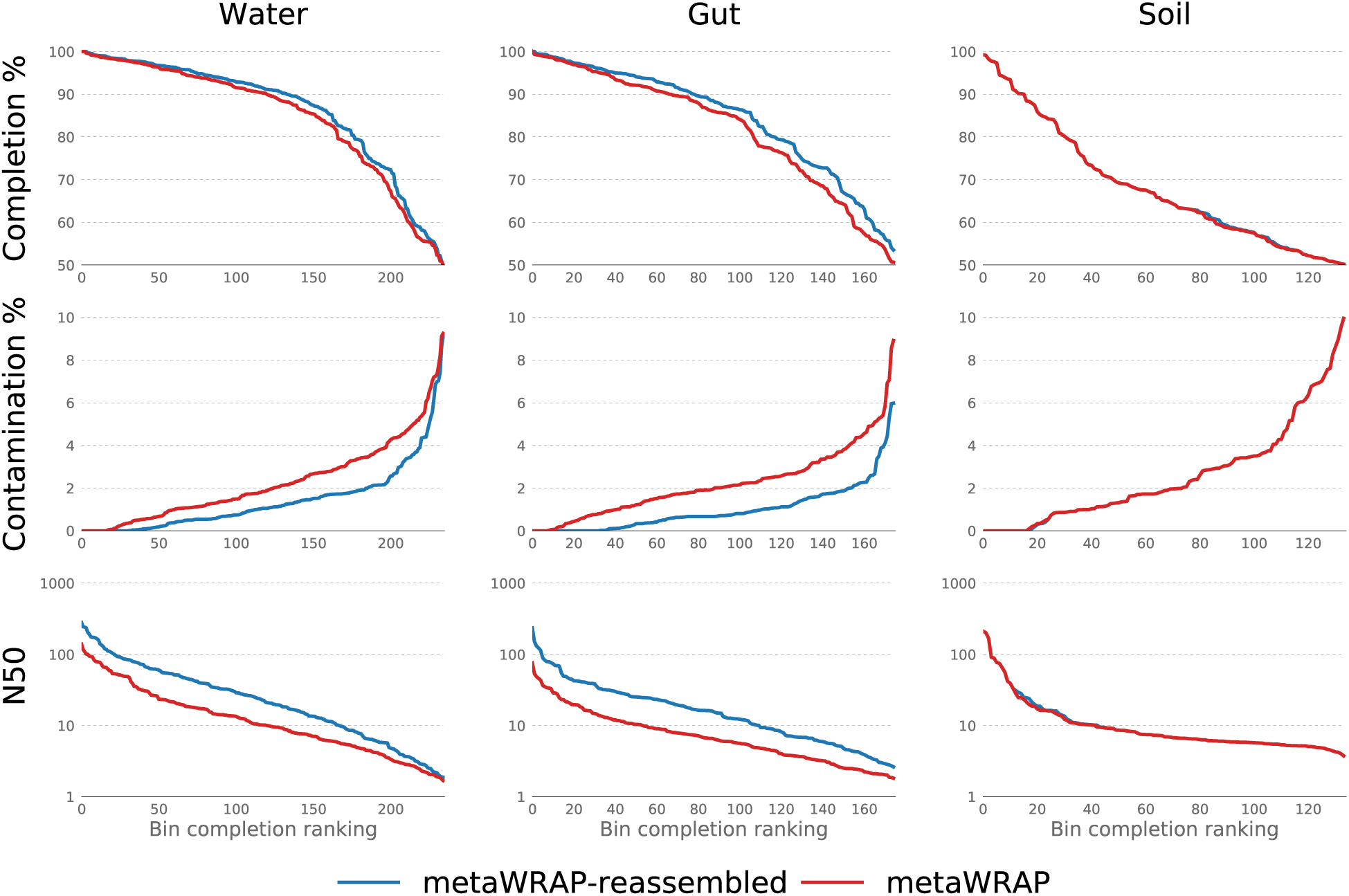
N50, completion, and contamination metrics of original bins extracted from water, gut, and soil metagenomes with the metaWRAP’s Bin_refinement module and the same bins reassembled with metaWRAP’s Reassemble_bins module. Only bins with≥50% completion and≤10% contamination are shown (estimated with CheckM).

### MetaWRAP produced high-quality draft genomes

We investigated the performance of different binning approaches (both original binners and bin consolidation software) when extracting high quality draft genomes, with a contamination less than 5% and completion greater than 70%, 80%, 90%, and 95%. The default run of metaWRAP-Bin_refinement consistently produced the highest number of high-quality draft genomes in water, gut, and soil data sets. These numbers further improved when re-running the module with appropriate minimum completion (-c) settings (i.e running Bin_refinement –c 90 when benchmarking for bins with a minimum completion of 90%). This approach significantly outperformed every other tested binning and bin refinement method at every quality threshold (Figure 6).

**Figure 6:**
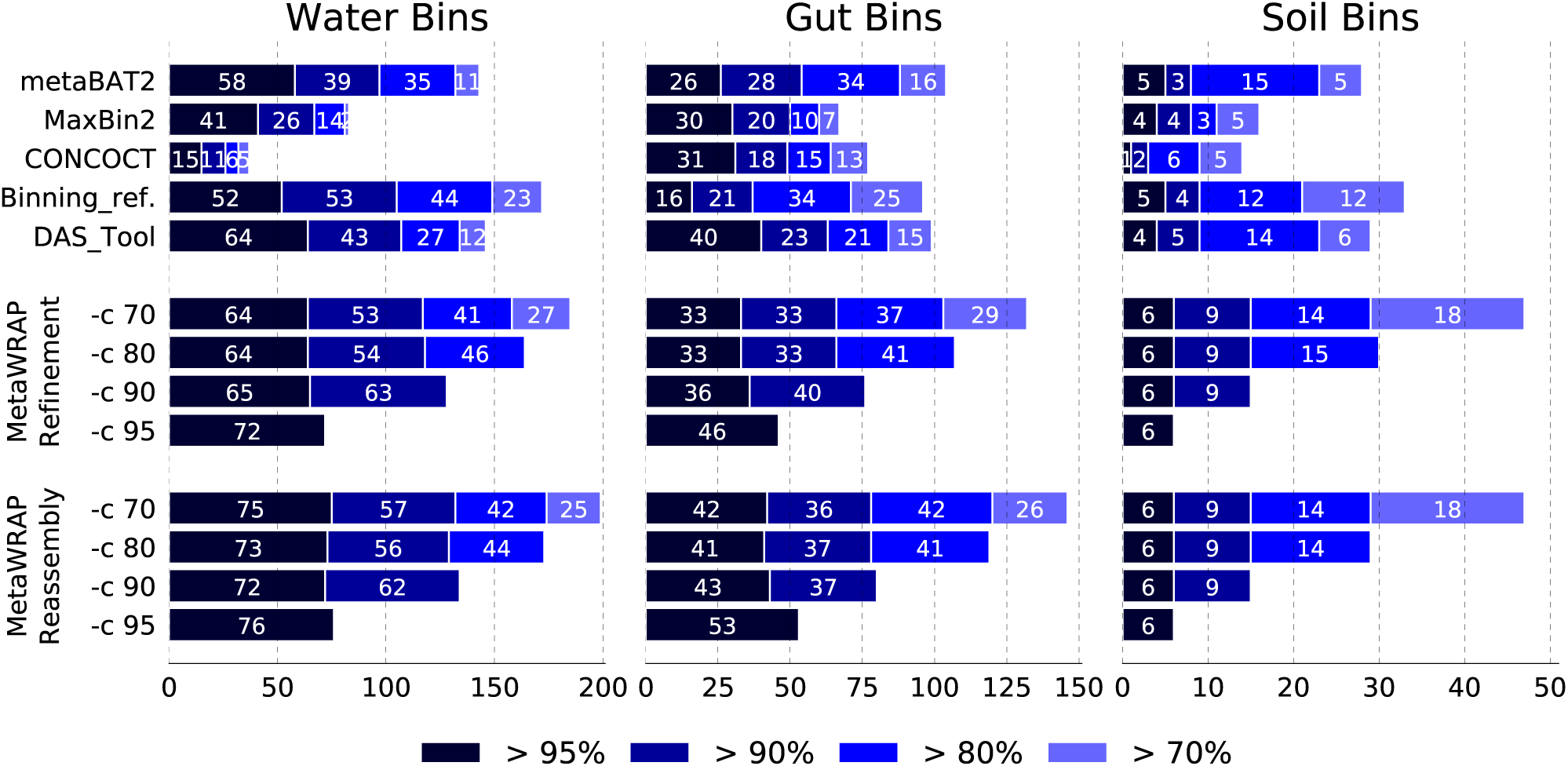
Number of high purity bins (less than 5% contamination) extracted from water, gut, and soil metagenomes with 70%, 80%, 90%, and 95% completion (estimated with CheckM) using original binning software (metaBAT2, MaxBin2, and CONCOCT) and bin refining algorithms (Binning_refiner, DAS_Tool, metaWRAP-Bin_refinement, and metaWRAP-Reassemble_bins). MetaWRAP modules were run with varying –c (minimum completion) parameters. MetaWRAP’s Reassemble_bins module was run on the output of the Bin_refinement module.

The reassembly of the metaWRAP-derived bins with the Reassemble_bins module made a further improvement on the number of high-quality draft genomes extracted from the gut and water data sets. Even the default run of Reassemble_bins produced a significantly better bin set compared to non-reassembled bin sets produced by all tested software, including metaWRAP’s Bin_refinement. However, just like in the Bin_refinement runs, the results were further enhanced when Reassemble_bins was provided with an appropriate –c option.

When comparing to the original binning software (MaxBin2, metaBAT2, and CONCOCT) and bin consolidation tools (DAS_Tool and Binning_refiner), metaWRAP produced the largest number of high-quality draft genomes in all the tested WMG data sets. Additionally, it should also be considered that metaWRAP is capable of improving bin sets from any binning software. Therefore, when new metagenomic binning software are developed, their outputs can still be used with metaWRAP refinement and reassembly algorithms.

### MetaWRAP enables analysis and visualization of metagenomic bins

The rest of metaWRAP modules address examining and processing a set of bins in preparation for downstream analysis. The user may visualize the bins in the context of the entire community with the Blobology module, quantify their abundances across samples with the Quant_bins module, estimate their taxonomy with the Classify_bins module, and functionally annotate them with the Annotate_bins module.

The metaWRAP-Quant_bins module was used to estimate bin abundances across samples from their respective microbiome survey, and the results were shown in a clustered heatmap (Additional file 11: Figure S8). Clustered heatmaps may be used to infer bin co-abundance and to identify similarities and differences between samples. Because this approach considers the abundances of every extracted bin individually, it offers higher resolution information than when using higher taxonomic ranks.

Bins were also visualized with the metaWRAP-Blobology module. The module produces GC vs Abundance plots of contigs, annotated with their taxonomy[43] (Figure 3), bin membership (Figure 7), or both (Additional file 12: Figure S9). These plots allow for inspection of the extracted bins in the context of the entire community that they belong to, as well as visualize the relative success of the binning process.

**Figure 7:**
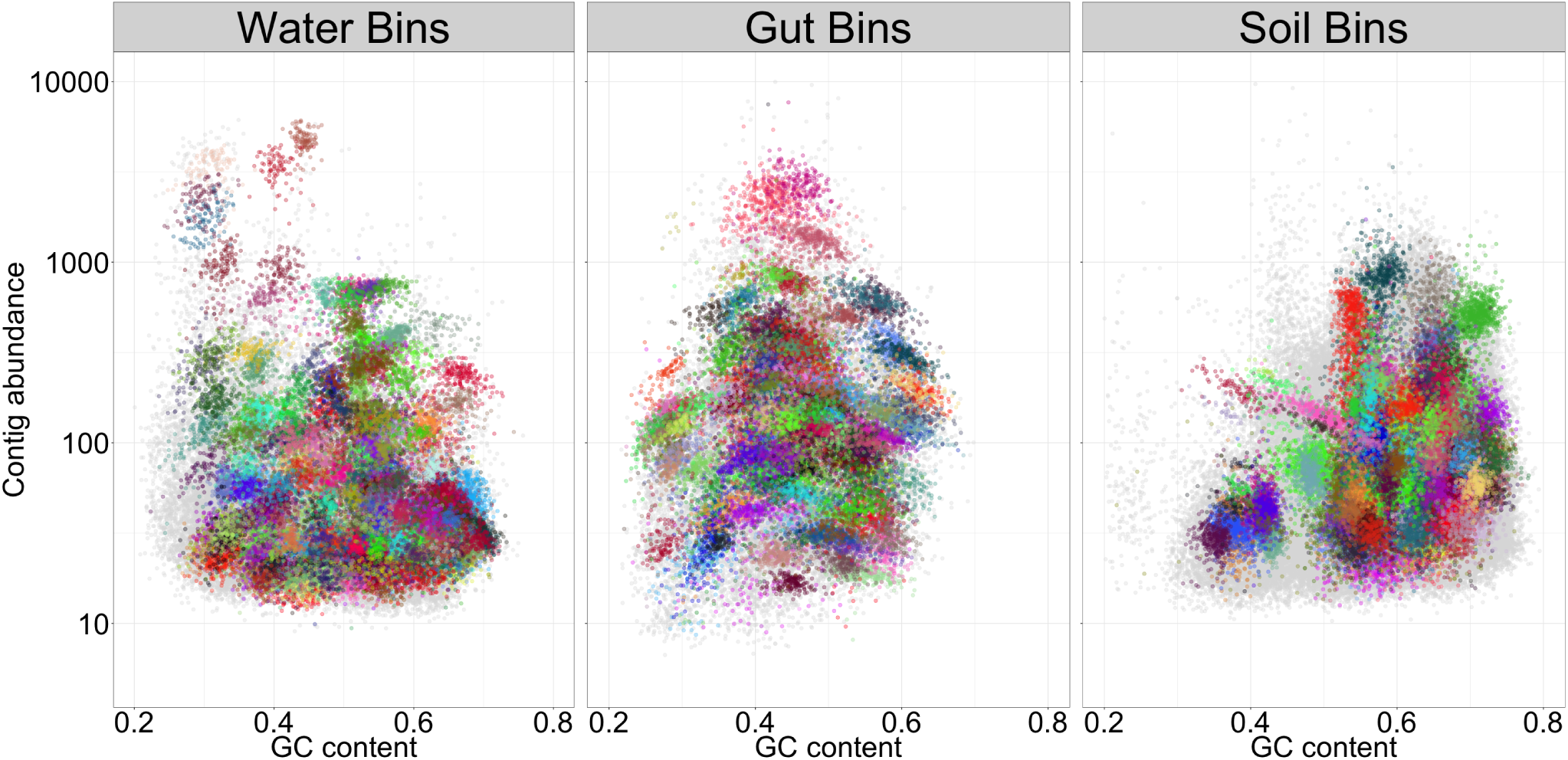
GC vs. abundance plots of contigs from water, gut, and soil metagenomes, produced with the Blobology module. Abundance of contigs was calculated from standardized read coverage in each sample. The contigs were annotated with the bins that they belong to (bin colors are chosen at random), allowing for quick inspection of binning success. Bins were produced with metaWRAP’s Bin_Refinement module. Only bins with ≥70% completion and≤10% contamination are shown (estimated with CheckM).

The final reassembled bins were taxonomy profiled with the metaWRAP-Classify_bins module (Additional file 13: Bin taxonomy) and functionally annotated with the Annotate_bins module. Together, this information may be used in downstream analysis to investigate complex questions about functional interactions and metabolic potential of individual community members.

## Conclusions

Genome-level analysis of WMG sequencing data is essential in understanding the composition and function of microbiomes. Until now, this rapidly growing field lacked a unifying platform to utilize the wealth of currently available software and make them easily accessible to researchers. MetaWRAP is a flexible pipeline that can handle common tasks in metagenomic data analysis starting from raw read quality control, and ending in bin extraction and analysis. MetaWRAP is easy to install through Bioconda, simple to use, and its modularity gives the investigator flexibility in their analysis approach.

MetaWRAP contributed significant improvements to the recovery of draft genomes from shotgun metagenomic data through bin refinement and reassembly. The bin refinement module uses a novel hybrid approach to consolidate bin predictions from different binning software, producing a single stronger set. This approach significantly outperformed individual binning software, as well as other consolidation algorithms. The algorithm can adjust to accommodate specific draft genome quality targets, making it suitable for many research applications. MetaWRAP’s bin reassembly module further improved the draft genomes in both completeness and purity. Finally, metaWRAP contains multiple modules for analysis and evaluation of metagenomic bins – bin taxonomy assignment, abundance estimation, functional annotation, and visualization.

## Methods

### CAMI binning benchmarking

The “gold standard” assemblies from the “high”, “medium”, and “low” diversity CAMI challenges were binned with metaBAT2 v2.12.1[22], Maxbin2 v2.2.4[21], and CONCOCT v0.4.0[19] using the metaWRAP-Binning module with default parameters. The resulting three bin sets were consolidated with DAS_Tool v1.1.0[28] (default settings, blast used for search engine), Binning_refiner v1.2[29] (default settings), and metaWRAP-Bin_refinement v0.7 (see Supp. Methods for module details). The completion and contamination of the bins in all six bin sets were first evaluated with CheckM v1.0.7[25] with default parameters, and bins with a completion less than 50% or a contamination greater than 10% were discarded. The true recall and precision of the bins within the six resulting bin sets was determined with Amber v0.6.2[40] and bin recall and precision were converted to completion and contamination percentages.

### Real data binning benchmarking

The raw sequences from water, gut, and soil microbiomes were run through the metaWRAP-Read_qc module, which trims the reads with TrimGalore[44], removes human contamination with BMTagger[45] searching againsts hg38, and produces a quality report with FASTQC[46]. MetaWRAP’s Kraken module (-s 10000000) was run on the quality-controlled with Kraken[30] (using standard database) and KronaTools 2.7[37]. The reads were co-assembled within each community type with MegaHit v1.1.2[35] by using the metaWRAP-Assembly module. Contigs shorter than 1000bp were discarded, with the exception of the soil assembly, for which the cutoff of 3000bp was chosen to reduce binning time. The co-assemblies of each data type were binned with metaBAT2 v2.12.1, Maxbin2 v2.2.4, and CONCOCT v0.4.0 using the metaWRAP-Binning module at default settings. The resulting three bin sets of each microbiome type were then passed to DAS_Tool v1.1.0 (–search_engine blast option), Binning_refiner v1.2 (default settings), and metaWRAP-Bin_refinement v0.7. For the main benchmark, metaWRAP was run with –c 50 –x 10 settings. To benchmark the bins produced by all the binning methods, the completion and contamination of the bins was estimated with CheckM v1.0.7. See Supplementary Methods for details on all modules.

### Bin_refinement optimization demonstration

The metaWRAP-Bin_Refinement module was run at different –c (minimum completion) and –x (maximum contamination) settings. First, the bin sets produced with metaBAT2 v2.12.1, Maxbin2 v2.2.4, and CONCOCT v0.4.0 were refined with the module with a constant maximum contamination setting –x 10, but varying minimum completion settings –c 50, 60, 70, 80, 90, and 95. Then the same bin sets were refined with a constant minimum contamination setting –c 50, but varying maximum contamination setting of –x 10, 8, 6, 4, 2, and 1.

### Reassembly benchmarking

Bin sets produced by the metaWRAP-Bin_refinement module with -c 50 –x 10 settings were run through the metaWRAP-Reassemble_bins (see Supp. Methods for module details) module with -c 50 –x 10 settings. The re-assembly module uses BWA 0.7.15[33] and Samtools 1.6[47] to pull reads belonging to each bin, and then reassembled them with SPAdes[42] (–carefull option). The resulting bins were evaluated with CheckM v1.0.7, and the completion and contamination values were sorted and plotted.

### Extracting high-quality draft genomes

MetaWRAP’s Bin_refinement and Reassemble_bins modules were run with contamination less than 5% and completion greater than 70%, 80%, 90%, or 95%. The Bin_refinement module was run on bin sets produced with metaBAT2 v2.12.1, Maxbin2 v2.2.4, and CONCOCT v0.4.0 with four different settings: -c 70 –x 5, -c 80 –x 5, -c 90 –x 5, -c 95 –x 5. The Reassemble_bins module was run with the same settings on the output of Bin_refinement with -c 60 –x 10, -c 70 –x 10, -c 80 –x 10, and –c 90 –x 10 settings, respectively. The resulting metaWRAP bin sets, the original bin sets, and the refinements from DAS_Tool and Binning_refiner were evaluated with CheckM v1.0.7 and the number of bins with different completion and contamination values were counted and plotted.

### Draft genomes analysis

Bins produced with metaWRAP-Bin_refinement (-c 70 –x 10 options) were visualized with the Blobology module (–bins flag used to provide bins), which uses a modified Blobology[38] scripts, Bowtie2 2.3.2[48], and MegaBLAST 2.6.0[43] to make Taxon-Annotated-GC-Coverage plots. Bin abundance in each sample was estimated and visualized with the Quant_bins module, which uses Salmon 0.9.1[49] to quantify individual contigs[50] and then estimate bin abundances. The reassembled bins from the metaWRAP-Reassemble_bins module (-c 50 –x 10 options) were run through the Classify_bins module (default settings), which makes initial taxonomy predictions of individual scaffolds with Taxator-tk 1.3.3e[31] and estimates the taxonomy of entire bins. Bins were functionally annotated with the metaWRAP-Annotate_bins module, which uses PROKKA 1.12[51] to annotate each bin in parallel.

## Availability and Requirements

**Project name:** metaWRAP

**Project home page:** https://github.com/bxlab/metaWRAP

**Operating system:** Linux64

**Programming languages:** Shell

**Other requirements:** Conda

**License:** MIT

## Abbreviations

WMG: whole metagenome
comp: completion
cont: contamination
-c: minimum completion parameter
-x: maximum contamination parameter

## Availability of data and materials

The datasets supporting the conclusions of this article are available from the original CAMI challenge (https://data.cami-challenge.org/participate) for the synthetic data sets, the National Center for Biotechnology Information under SRA numbers SRR2053273–SRR2053308 for the Central Baltic Surface Water Metagenome, SRA numbers ERR011087-ERR011136 for the Metagenomic of the Human Intestinal Tract (MetaHIT) survey, and at Joint Genome Institute under Gold Analysis Project IDs Ga0007435, Ga0007436, Ga0007437, Ga0007438, Ga0007439, and Ga0007440 for the soil data. All analysis results and scripts used to generate figures are available at https://github.com/ursky/metawrap_paper.

## Author’s contributions

GU built, released, and maintained the metaWRAP software, ran the benchmarks, and wrote the manuscript. JDR and JT provided ideas for building and improving metaWRAP and edited the manuscript. All authors read and approved the final manuscript.

## Acknowledgements

We thank early users of metaWRAP Alejandro Palomo, Keith Arora-Williams, and Emine Ertekin for their patience and help with debugging, Bing Ma for module suggestions, and Michael Sauria for computational support.

---

**Figure S1:**
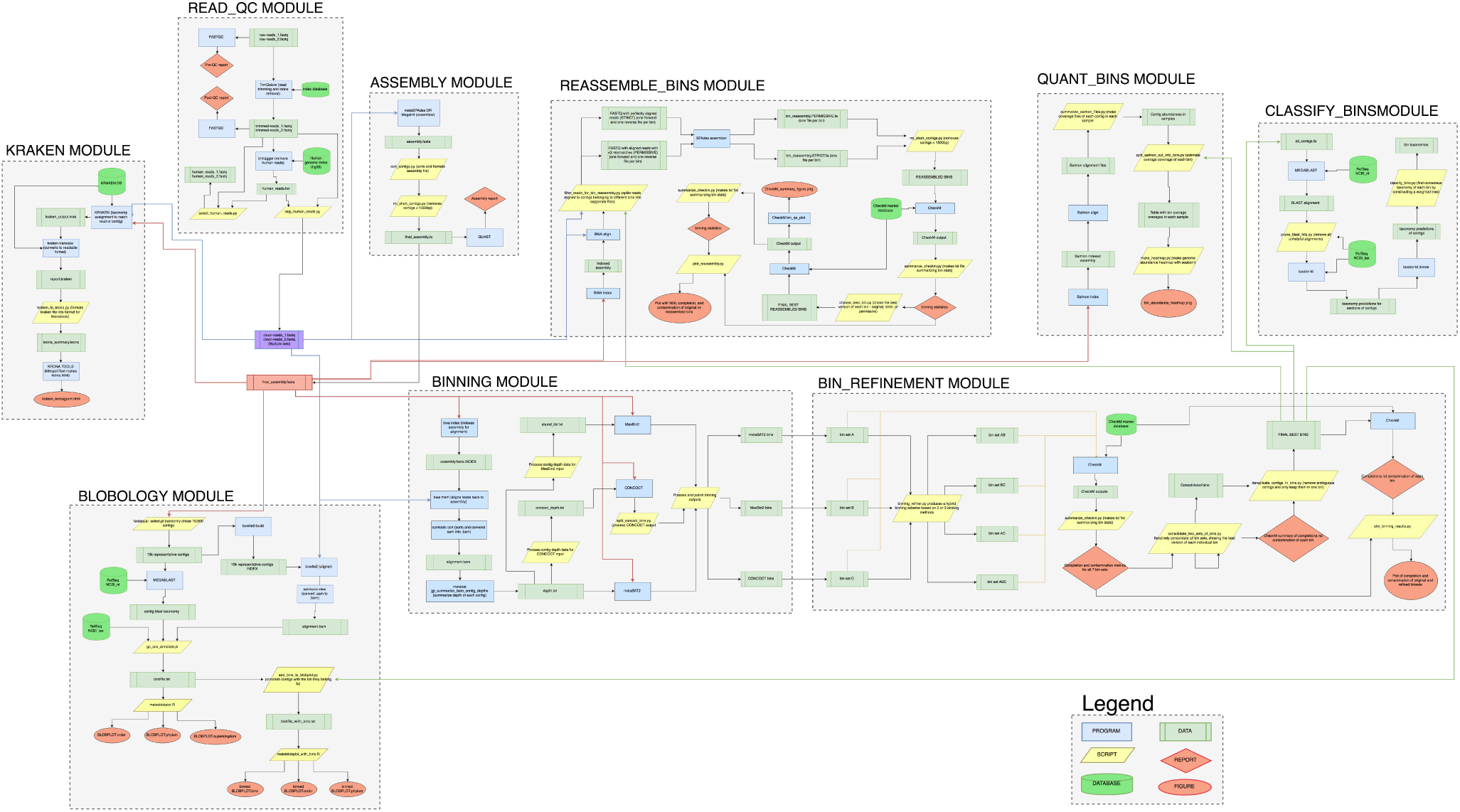
Detailed walkthrough of the data files, software, databases, and custom scripts that metaWRAP uses. The components of each metaWRAP module grouped and denoted with dotted lines.

**Figure S2:**
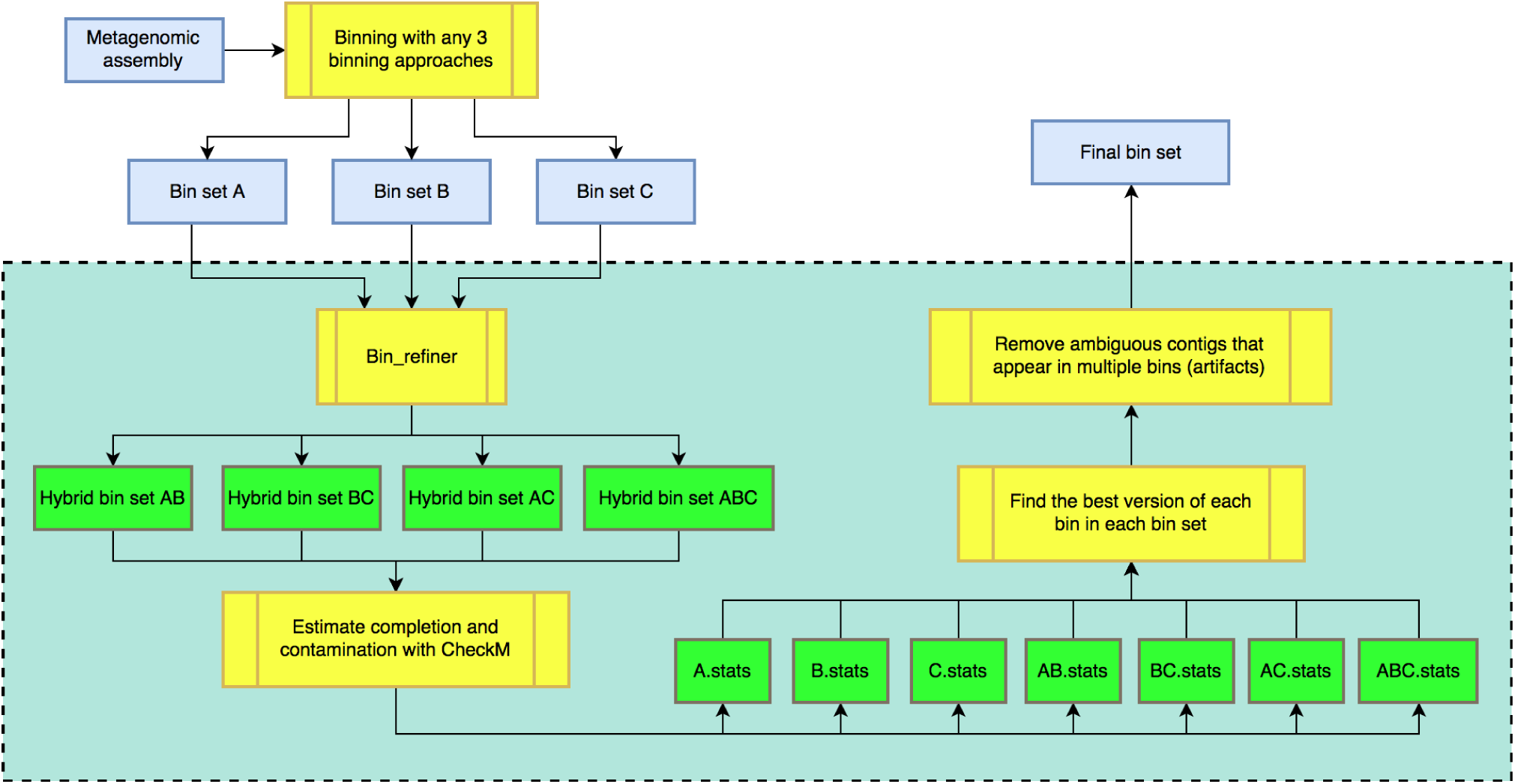
Logical workflow of the Bin_refinement modules of metaWRAP. The module takes in three bin sets produced from the same assembly by different software or different parameters of the same software. Binning_refiner is used to create hybridized intermediates (4 possible combinations), and the completion and contamination of the original and hybridized bins is estimated with CheckM. The best version of each bin is then found in the resulting 7 bin sets.

**Figure S3:**
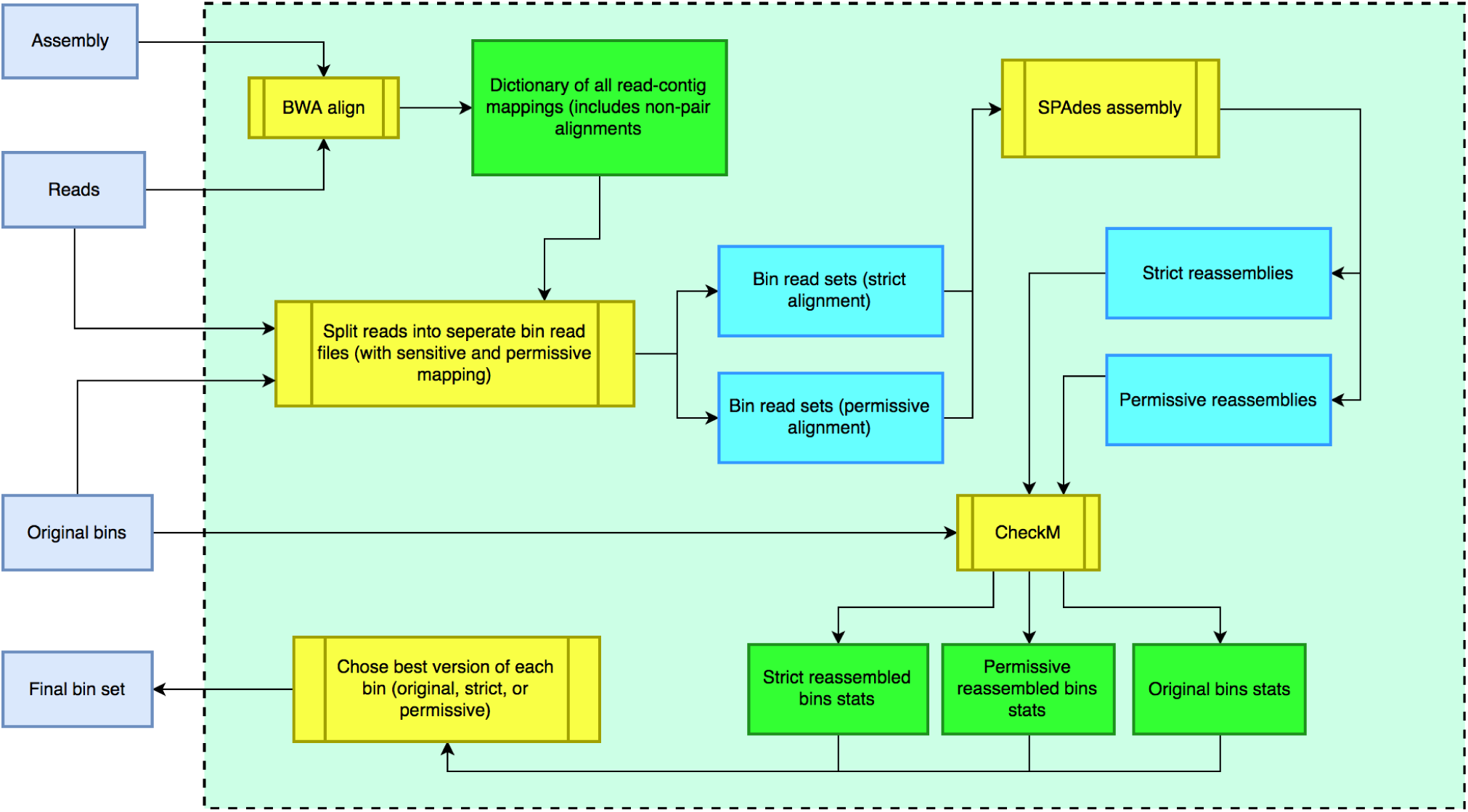
Logical workflow of the Reassemble_bins module, which extracts reads belonging to bins in a given bin set, and individually reassembles them. This process is done for perfectly mapping reads (strict) and reads mapping with less than 3 mismatches (permissive). For each bin, CheckM is used choose the best bin between the original and the two reassembled versions.

**Figure S4:**
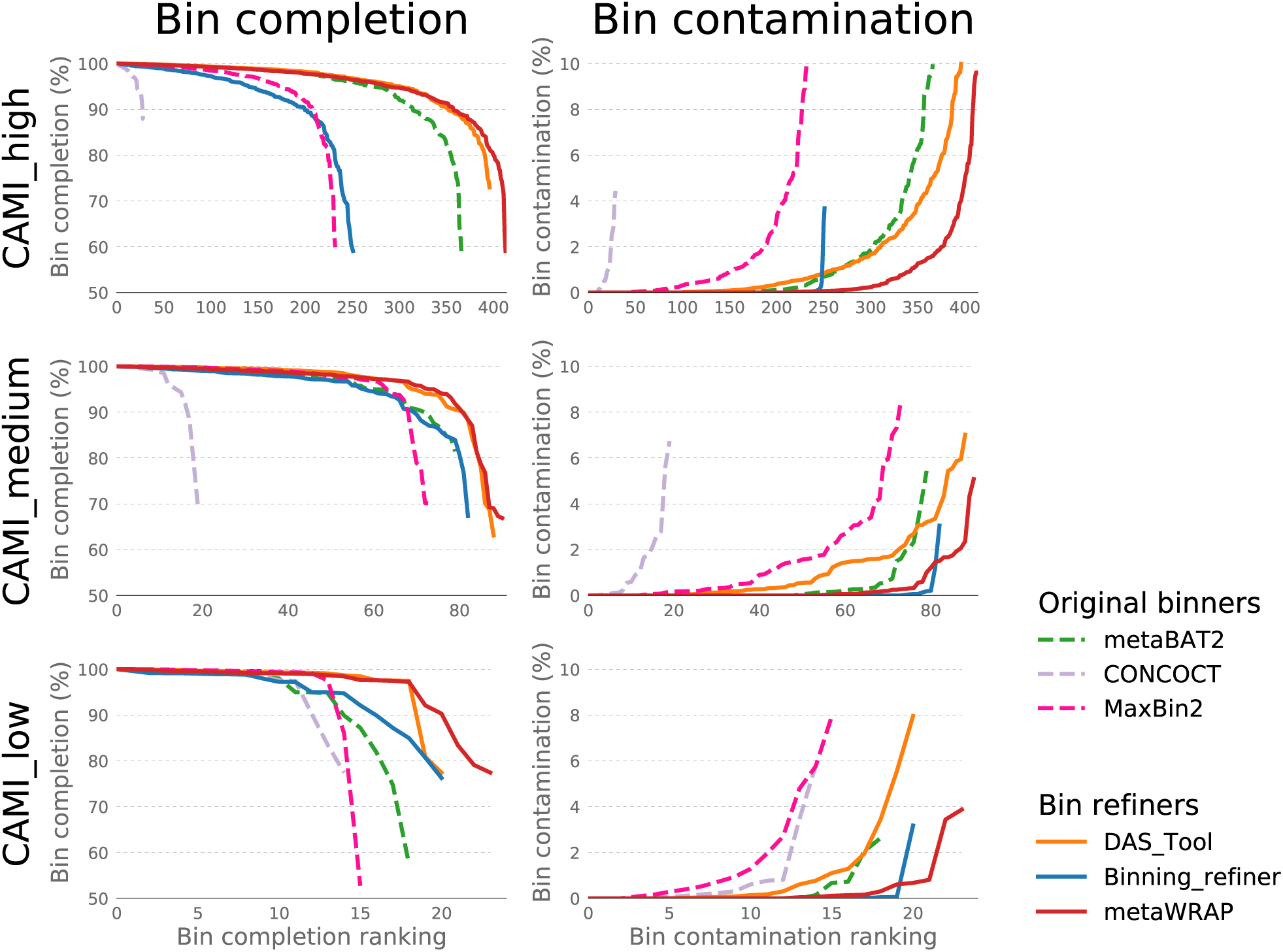
Completion and contamination (determined with CheckM) of bins recovered from the CAMI’s high, medium, and low complexity synthetic data sets using original binning software (metaBAT2, MaxBin2, CONCOCT) and software consolidating the original sets (DAS_Tool, Binning_refiner, metaWRAP). Only bins with≥50% completion and≤10% contamination are shown.

**Figure S5:**
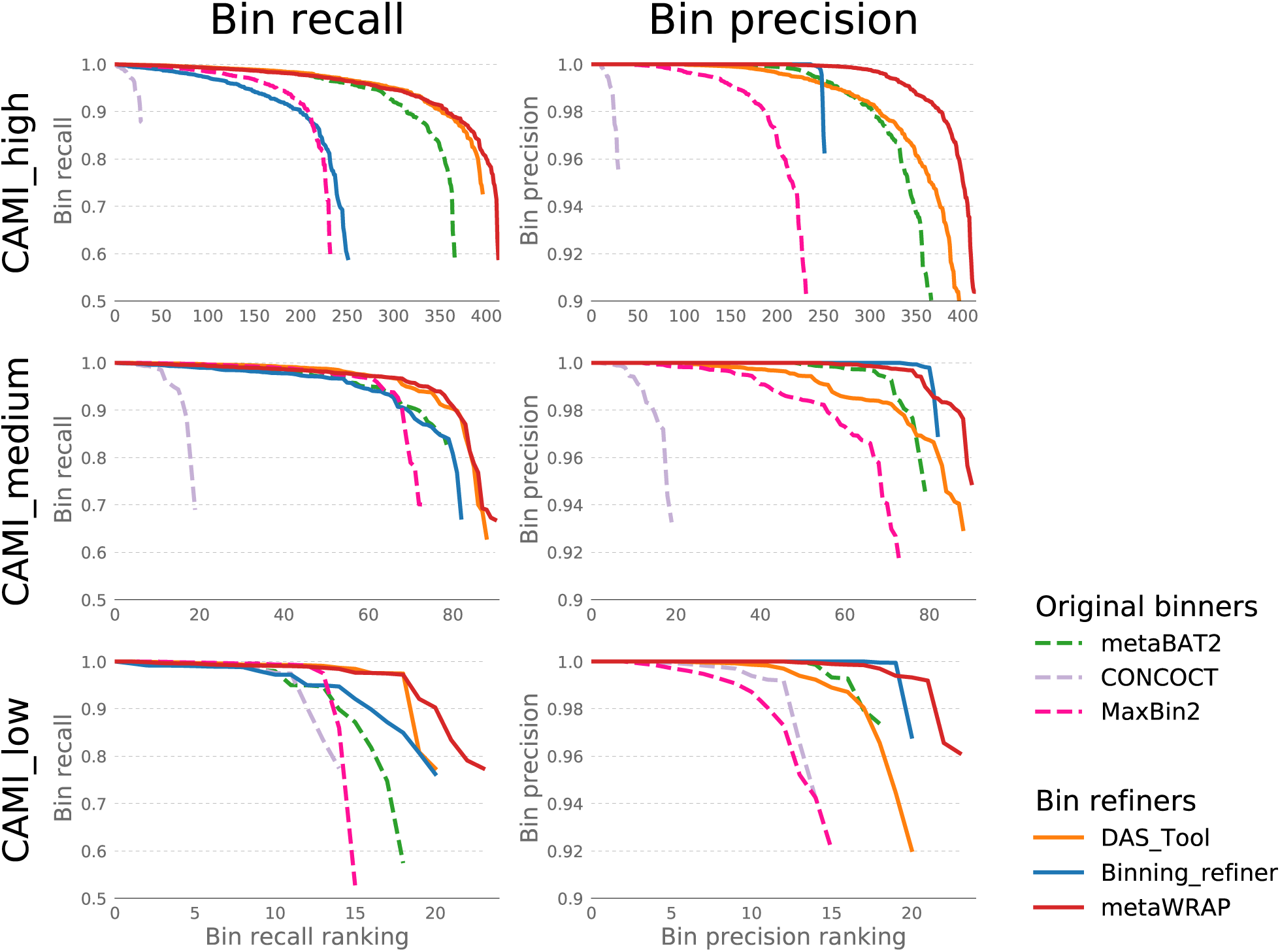
True recall and precision (determined with AMBER) of bins recovered from the CAMI’s high, medium, and low complexity synthetic data sets using original binning software (metaBAT2, MaxBin2, CONCOCT) and software consolidating the original sets (DAS_Tool, Binning_refiner, metaWRAP). Only bins with ≥0.5% recall and ≥0.9% precision are shown.

**Figure S6:**
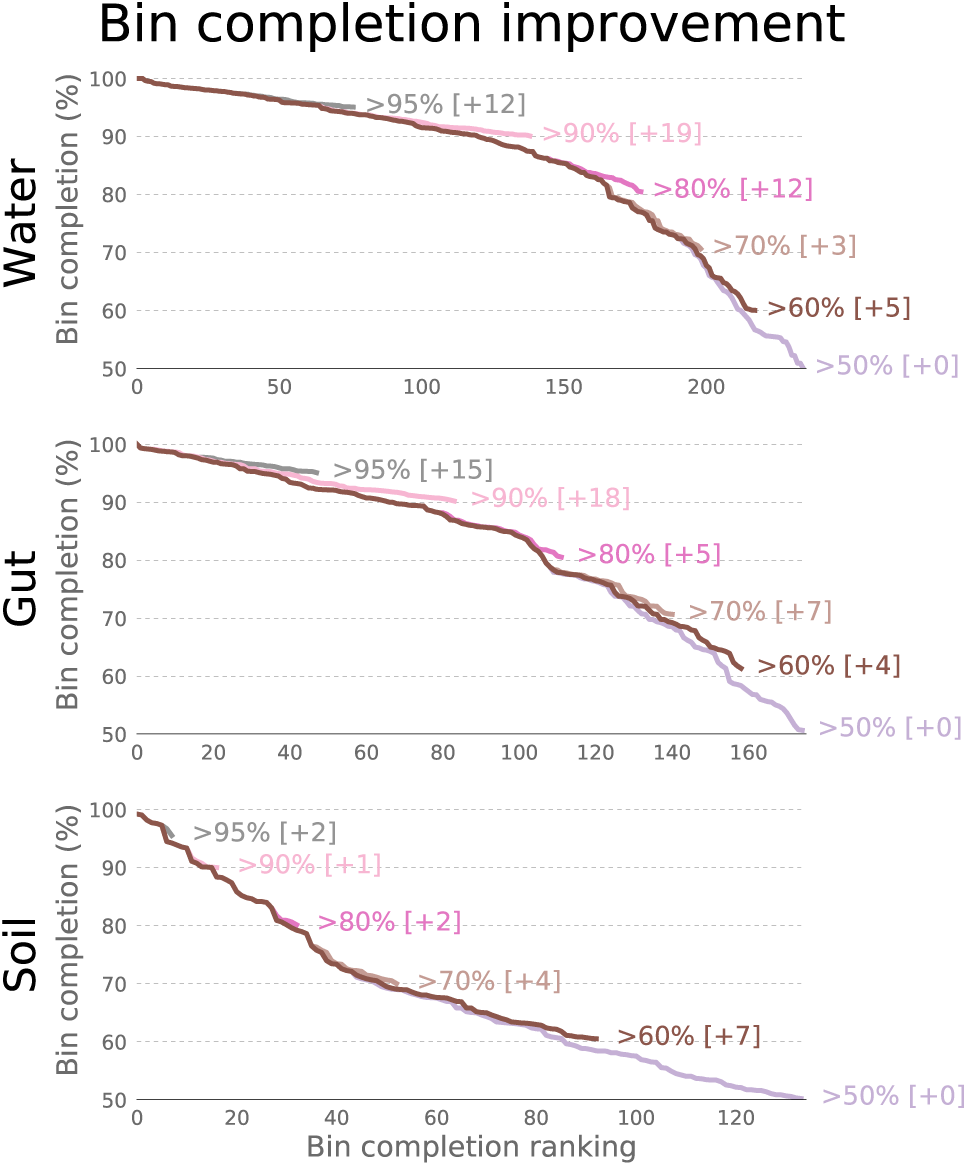
Completion of bins recovered from water, gut, and soil metagenomes with the metaWRAP Bin_refinement module with a varying minimum completion parameter (-c), but constant maximum contamination parameter (-x 10). The numbers in the brackets indicate the number of extra bins gained at that threshold compared to the baseline run (-c 50 –x 10). Only bins with ≥50% completion and ≤10% contamination are shown.

**Figure S7:**
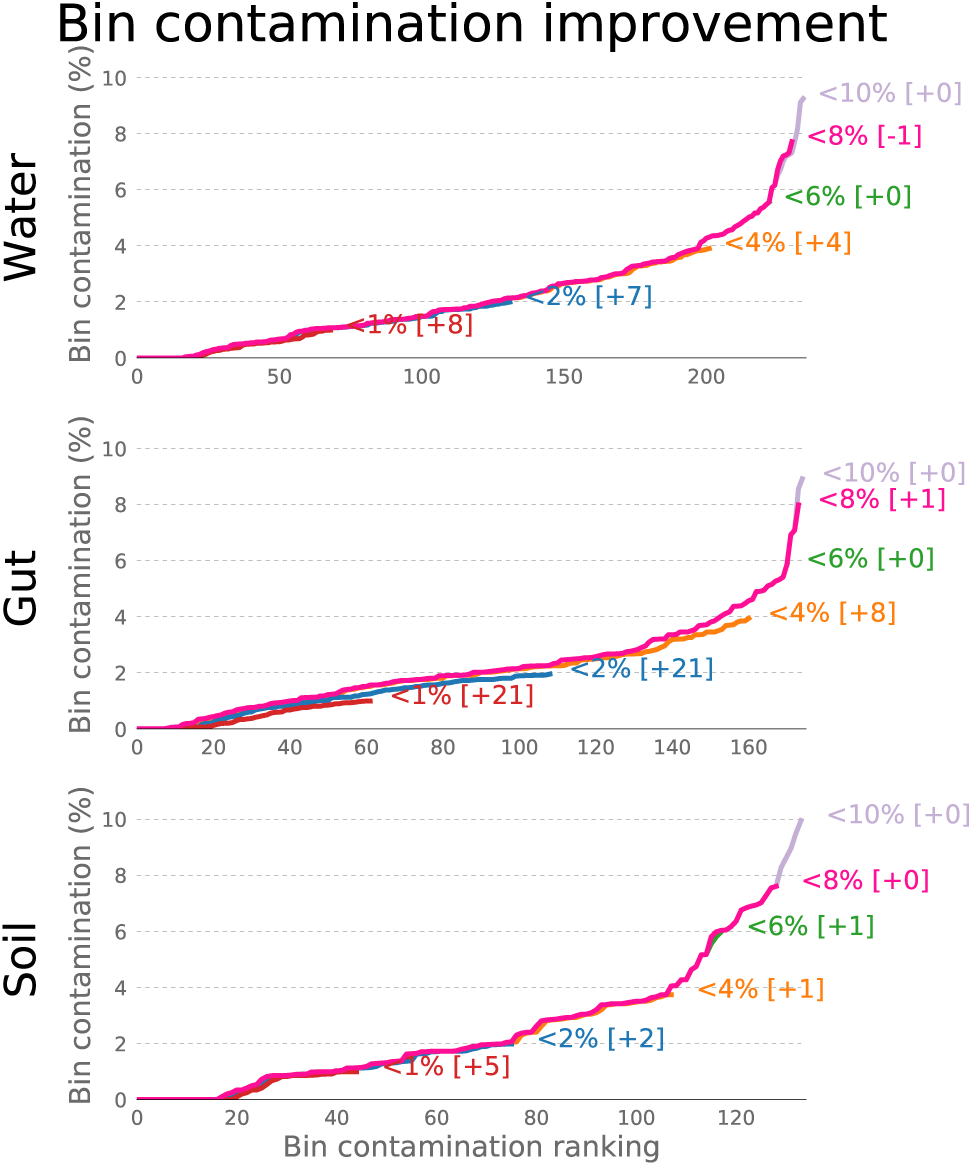
Contamination of bins recovered from water, gut, and soil metagenomes with the metaWRAP Bin_refinement module with a varying maximum contamination parameter (-x), but constant minimum completion parameter (-c 50). The numbers in the brackets indicate the number of extra bins gained at that threshold compared to the baseline run (-c 50 –x 10). Only bins with≥50% completion and≤10% contamination are shown.

**Figure S8:**
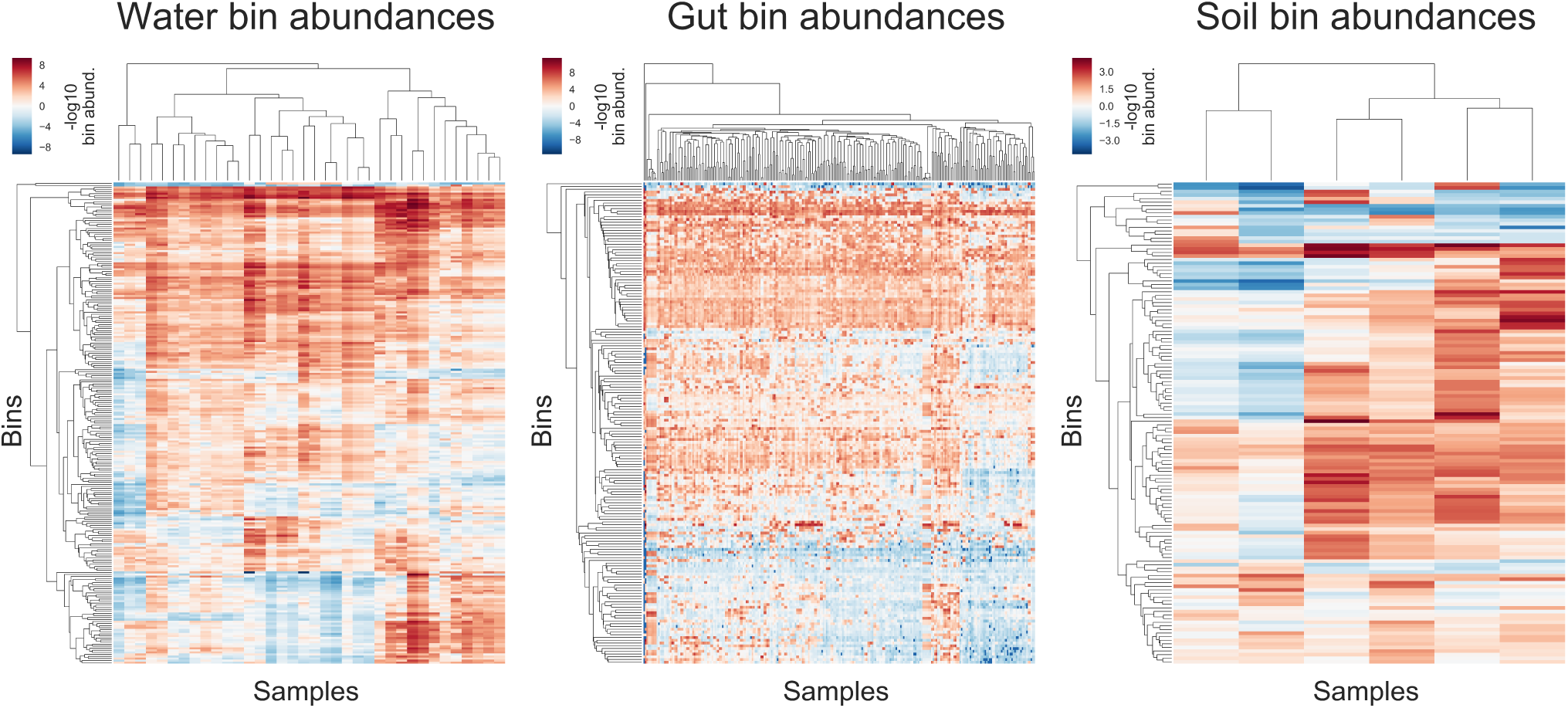
Clustered heat maps showing the log of bin abundance of bins extracted with metaWRAP Bin_refinement (-c 50 –x 10) across samples in water, gut, and soil metagenomes, calculated and plotted with metaWRAP’s Quant_bins module.

**Figure S9:**
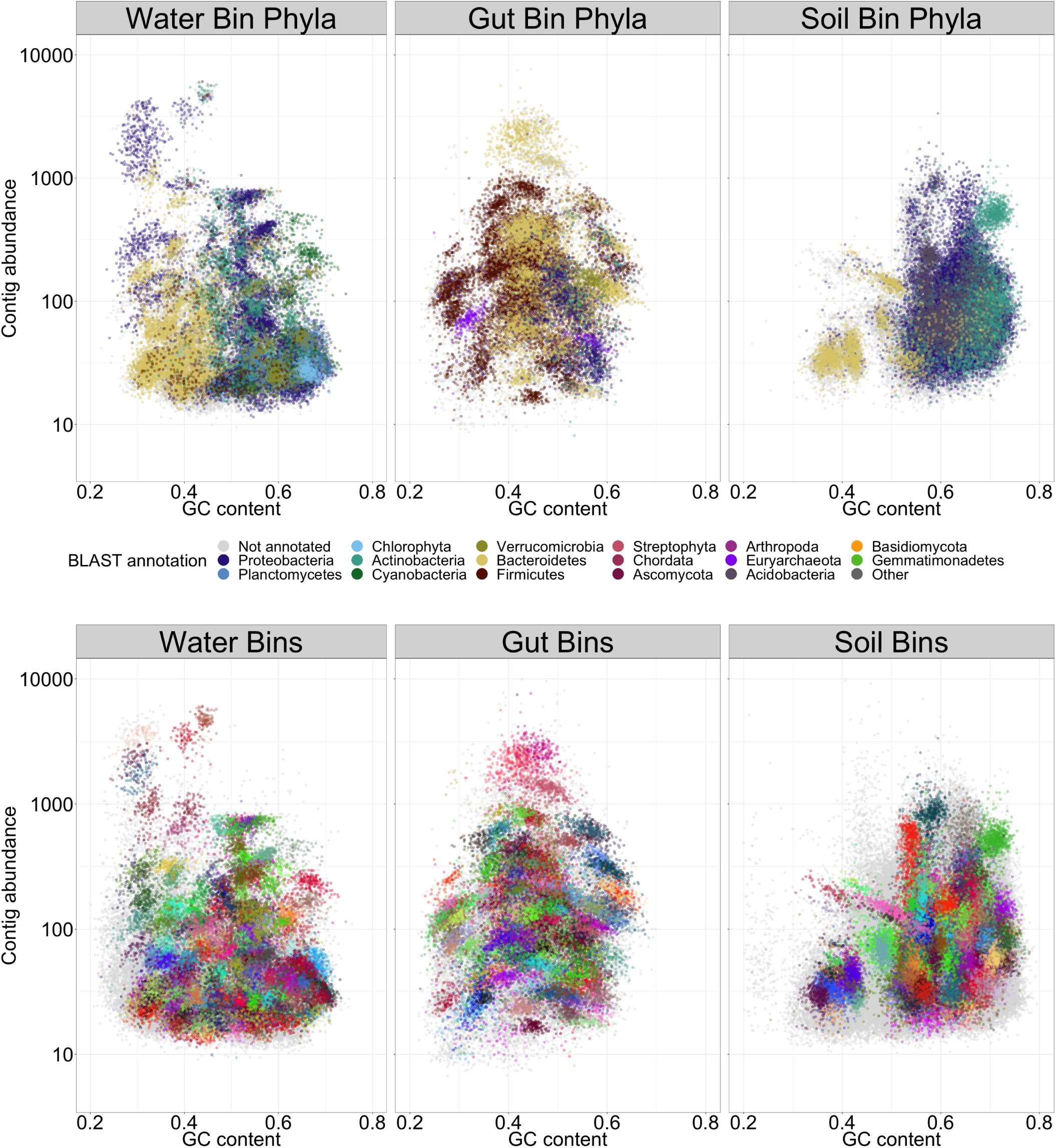
MetaWRAP’s Blobology module visualization of water, gut, and soil metagenomes, showing the GC and average coverage of each successfully binned contig (metaWRAP Bin_refinement –c 70 –x 10) in the assemblies, and annotated with the taxonomy at the phylum level, and the bins that they belong to (bin colors are chosen at random).

